## Supplemental Material for "Working Memory Gating in Obesity is Moderated by Striatal Dopaminergic Gene Variants"

**SUPPLEMENTARY MATERIALS´**

**Model comparisons**

Model 1:
model_BMI_reduced:
correct ~ condition * zBMI + zIQ + zWM_tired + zWM_conc + Gender + (1 | ID)

model_BMI_full:
correct ~ condition * zBMI + zIQ + zWM_tired + zWM_conc + Gender + BED + project + zDFS + zAge + (1 | ID)

| Model | N. of Parameters | AIC | BIC | Log Likelihood | Deviance | Chi-Squared | Df | p-value |
|---|---|---|---|---|---|---|---|---|
| model reduced | 13 | 23060 | 23172 | -11517 | 23034 | | | |
| model_full | 18 | 23068 | 23222 | -11516 | 23032 | 2.41 | 5 | 0.791 |

Model 2:
model_COMT_Taq1A_BMI_reduced:
correct ~ condition * COMT * DRD2 * zBMI + zIQ + Gender + zWM_tired + zWM_conc + (1 | ID)

model_COMT_Taq1A_BMI_full:
correct ~ condition * COMT * DRD2 * zBMI + zIQ + Gender + zWM_tired + zWM_conc + zAge + project + BED + zDFS + (1 | ID)

| Model | N. of Parameters | AIC | BIC | Log Likelihood | Deviance | Chi-Squared | Df | p-value |
|---|---|---|---|---|---|---|---|---|
| model reduced | 53 | 23091 | 23545 | -11492 | 22985 | | | |
| model_full | 58 | 23098 | 23595 | -11491 | 22982 | 2.92 | 5 | 0.712 |

Model 3:
model_DARPP_BMI_reduced:
correct ~ DARPP*zBMI*condition + zIQ + zWM_conc + zWM_tired + Gender + (1 | ID)

model_DARPP_BMI_full:
correct ~ DARPP*zBMI*condition + zIQ + zWM_conc + zWM_tired + Gender + zAge + project + BED + zDFS + (1 | ID)

| Model | N. of Parameters | AIC | BIC | Log Likelihood | Deviance | Chi-Squared | Df | p-value |
|---|---|---|---|---|---|---|---|---|
| model reduced | 21 | 23055 | 23235 | -11506 | 23013 | | | |
| model_full | 26 | 23063 | 23286 | -11505 | 23011 | 2.39 | 5 | 0.793 |

Model 4:
model_C957T_BMI_reduced:
correct ~ C957T*zBMI*condition + zIQ + zWM_conc + zWM_tired + Gender + (1 | ID)

model_C957T_BMI_full:
correct ~ C957T*zBMI*condition + zIQ + zWM_conc + zWM_tired + Gender + zAge + project + BED + zDFS + (1 | ID)

| Model | N. of Parameters | AIC | BIC | Log Likelihood | Deviance | Chi-Squared | Df | p-value |
|---|---|---|---|---|---|---|---|---|
| model reduced | 21 | 23069 | 23249 | -11514 | 23027 | | | |
| model_full | 26 | 23077 | 23300 | -11512 | 23025 | 2.36 | 5 | 0.796 |

Model 5:
model_AA_BMI_reduced:
correct ~ scale(AAratio)*zBMI*condition + zIQ + zWM_conc + Gender + (1 | ID)

model_AA_BMI_full:
correct ~ scale(AAratio)*zBMI*condition + zIQ + zWM_conc + Gender + zWM_tired + zDFS + project + zAge + (1 | ID)

| Model | N. of Parameters | AIC | BIC | Log Likelihood | Deviance | Chi-Squared | Df | p-value |
|---|---|---|---|---|---|---|---|---|
| model reduced | 20 | 10812 | 10970 | -5386 | 10772 | | | |
| model_full | 24 | 10815 | 11005 | -5383 | 10767 | 5.06 | 4 | 0.281 |

**Check relationship amino acid ratio x condition x BMI for outlier**

Because there was an extreme BMI data point, we re-ran the model excluding this data point to check whether the results still hold. The three-way interaction between Amino Acid Ratio, BMI, and condition became trend-significant ($p_{corrected}$ = 0.063). It should be noted that, although this BMI data-point is a statistical outlier, it can still be considered as a valid data-point and might represent relevant variance in out data.

**Table S1**. Full output model 5 without extreme data point

| | Chisq | Df | Pr(>Chisq) |
|---|---|---|---|
| (Intercept) | 10.62 | 1 | 0.001 |
| AAratio | 0.54 | 1 | 0.461 |
| zBMI | 0.80 | 1 | 0.370 |
| condition | 2.98 | 3 | 0.394 |
| zIQ | 4.88 | 1 | 0.027 |
| zWM_conc | 11.55 | 1 | < .001 |
| Gender | 18.49 | 1 | < .001 |
| AAratio:zBMI | 0.28 | 1 | 0.597 |
| AAratio:condition | 7.28 | 3 | 0.064 |
| BMI:condition | 8.61 | 3 | 0.035 |
| AAratio:BMI:condition | 10.34 | 3 | 0.016 |

N = 159

Marginal $R^2$ / Conditional $R^2$ = 0.068 / 0.170

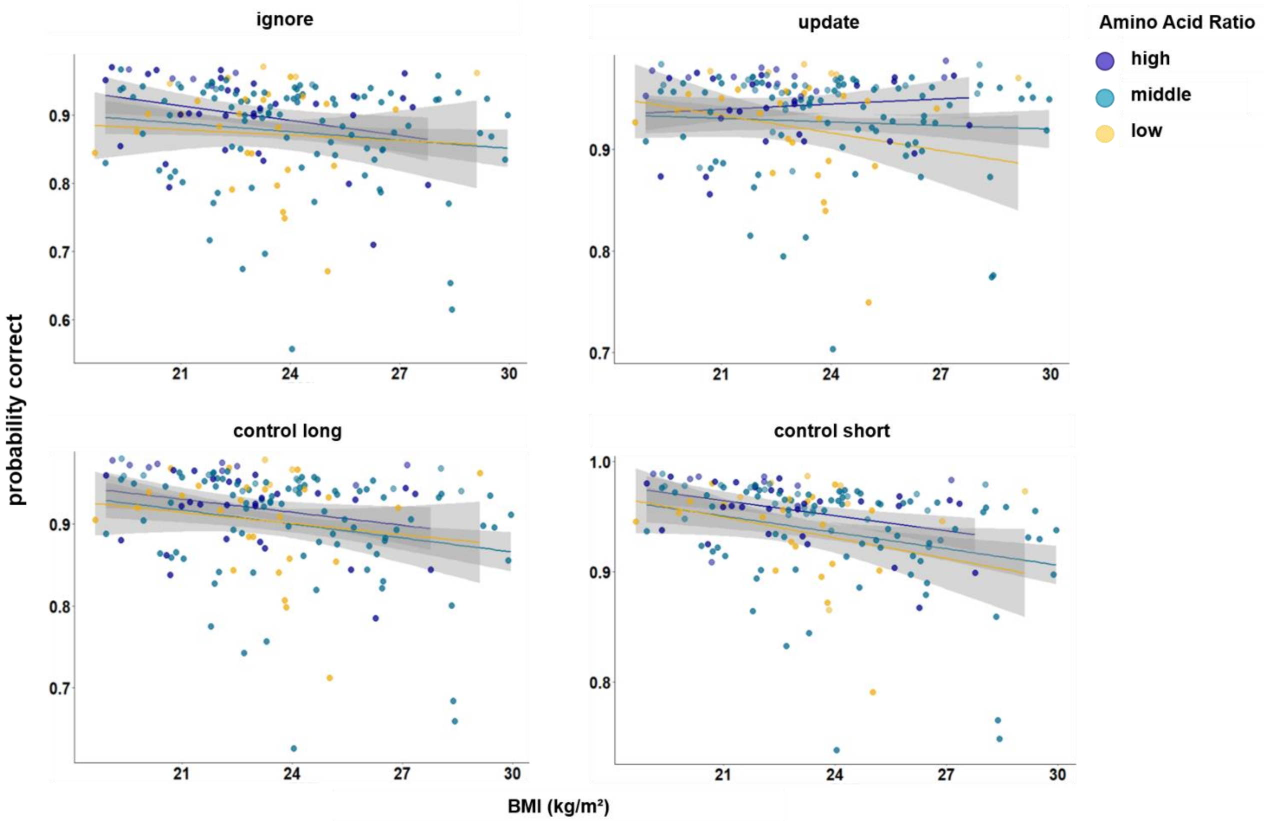


**Figure S1.** Interaction of Amino Acid Ratio, BMI and condition on data without the statistical BMI-outlier. Shaded areas represent the 95% confidence intervals.

**Interaction effects of various SNPs and Amino Acid Ratio**

Because it has been reported that there might be C957T and Taq1A interaction effects on working memory (Frank & Hutchison, 2009), we also looked at this interaction. Table S2 shows the full model output for the model investigating the interaction effect of C957T and Taq1A on accuracy in working memory conditions, depending on BMI.

Furthermore, we ran a model where we investigated the interaction of C957T and COMT, as also these two SNPs have been shown to interact (Xu et al., 2007). See Table S3 for the full model output.

Last but not least, since our initial analyses showed significant effects, we also looked at how amino acid ratio and Taq1A, and amino acid and DARPP would interact. We did not include all three factors in one model but ran separate models for both 4-way interactions since adding all factors would lead to uninterpretable interactions. Table S4 shows the full model output for the model investigating the interaction effect of Amino Acid Ratio and Taq1A. Table S5 shows the full model output for the model investigating the interaction effect of Amino Acid Ratio and DARPP. Both models looked at the effect of these interactions on accuracy in working memory conditions, depending on BMI.

**Table S2.** Full output for the model investigating C957T and Taq1A interaction effects

| | Chisq | Df | Pr(>Chisq) |
|---|---|---|---|
| (Intercept) | 276 | 1 | < .001 |
| condition | 26.9 | 3 | < .001 |
| C957T | 0.078 | 1 | 0.780 |
| Taq1A | 0.007 | 1 | 0.932 |
| zBMI | 6.47 | 1 | 0.011 |
| zIQ | 27.8 | 1 | < .001 |
| Gender | 7.75 | 1 | 0.005 |
| zWM_tired | 11.4 | 1 | < .001 |
| zWM_conc | 30.9 | 1 | < .001 |
| condition:C957T | 3.64 | 3 | 0.303 |
| condition:Taq1A | 2.97 | 3 | 0.396 |
| C957T:Taq1A | 0.047 | 1 | 0.828 |
| condition:zBMI | 1.21 | 3 | 0.751 |
| C957T:zBMI | 1.48 | 1 | 0.224 |
| Taq1A:zBMI | 2.21 | 1 | 0.138 |
| condition:C957T:Taq1A | 2.69 | 3 | 0.442 |
| condition:C957T:zBMI | 0.848 | 3 | 0.838 |
| condition:Taq1A:zBMI | 1.06 | 3 | 0.786 |
| C957T:Taq1A:zBMI | 0.596 | 1 | 0.440 |
| condition:C957T:Taq1A:zBMI | 1.8 | 3 | 0.615 |

N = 318

Marginal $R^2$ / Conditional $R^2$ = 0.074 / 0.172

**Table S3.** Full output for the model investigating C957T and COMT interaction effects

| | Chisq | Df | Pr(>Chisq) |
|---|---|---|---|
| (Intercept) | 265 | 1 | < .001 |
| condition | 34.9 | 3 | < .001 |
| C957T | 0.797 | 1 | 0.372 |
| COMT | 2.56 | 2 | 0.278 |
| zBMI | 2.62 | 1 | 0.106 |
| zIQ | 24.6 | 1 | < .001 |
| Gender | 8.73 | 1 | 0.003 |
| zWM_tired | 10.4 | 1 | 0.001 |
| zWM_conc | 32.6 | 1 | < .001 |
| condition:C957T | 4.33 | 3 | 0.228 |
| condition:COMT | 2.91 | 6 | 0.820 |
| C957T:COMT | 3.12 | 2 | 0.210 |
| condition:zBMI | 1.67 | 3 | 0.644 |
| C957T:zBMI | 0.113 | 1 | 0.736 |
| COMT:zBMI | 1.37 | 2 | 0.504 |
| condition:C957T:COMT | 1.38 | 6 | 0.967 |
| condition:C957T:zBMI | 1.45 | 3 | 0.694 |
| condition:COMT:zBMI | 15.2 | 6 | 0.019 |
| C957T:COMT:zBMI | 1.27 | 2 | 0.529 |
| condition:C957T:COMT:zBMI | 14.4 | 6 | 0.026 |

N = 318

Marginal $R^2$ / Conditional $R^2$ = 0.073 / 0.173

**Table S4.** Full output for the model investigating Amino Acid Ratio and Taq1A interaction effects

| | Chisq | Df | Pr(>Chisq) |
|---|---|---|---|
| (Intercept) | 18.5 | 1 | < .001 |
| condition | 2.81 | 3 | 0.422 |
| AAratio | 0.044 | 1 | 0.835 |
| Taq1A | 0.294 | 1 | 0.588 |
| zBMI | 1.75 | 1 | 0.185 |
| zIQ | 11.1 | 1 | < .001 |
| Gender | 4.9 | 1 | 0.027 |
| zWM_tired | 2.6 | 1 | 0.107 |
| zWM_conc | 22.1 | 1 | < .001 |
| condition:AAratio | 7.31 | 3 | 0.063 |
| condition:Taq1A | 0.743 | 3 | 0.863 |
| AAratio:Taq1A | 0.42 | 1 | 0.517 |
| condition:zBMI | 6.95 | 3 | 0.074 |
| AAratio:zBMI | 0.403 | 1 | 0.525 |
| Taq1A:zBMI | 0.174 | 1 | 0.676 |
| condition:AAratio:Taq1A | 0.784 | 3 | 0.853 |
| condition:AAratio:zBMI | 9.79 | 3 | 0.020 |
| condition:Taq1A:zBMI | 1.93 | 3 | 0.588 |
| AAratio:Taq1A:zBMI | 0.001 | 1 | 0.969 |
| condition:AAratio:Taq1A:zBMI | 2.07 | 3 | 0.559 |

N = 160

Marginal $R^2$ / Conditional $R^2$ = 0.077 / 0.172

**Table S5.** Full output for the model investigating Amino Acid Ratio and DARPP interaction effects

| | Chisq | Df | Pr(>Chisq) |
|---|---|---|---|
| (Intercept) | 20 | 1 | < .001 |
| condition | 6.92 | 3 | 0.075 |
| AAratio | 0.001 | 1 | 0.972 |
| DARPP | 0.147 | 1 | 0.701 |
| zBMI | 1.06 | 1 | 0.303 |
| zIQ | 10.1 | 1 | 0.001 |
| Gender | 3.52 | 1 | 0.061 |
| zWM_tired | 4.38 | 1 | 0.036 |
| zWM_conc | 22.6 | 1 | < .001 |
| condition:AAratio | 13.2 | 3 | 0.004 |
| condition:DARPP | 2.32 | 3 | 0.508 |
| AAratio:DARPP | 0.292 | 1 | 0.589 |
| condition:zBMI | 16.6 | 3 | < .001 |
| AAratio:zBMI | 0.273 | 1 | 0.601 |
| DARPP:zBMI | 0.017 | 1 | 0.896 |
| condition:AAratio:DARPP | 2.22 | 3 | 0.528 |
| condition:AAratio:zBMI | 17.2 | 3 | < .001 |
| condition:DARPP:zBMI | 7.4 | 3 | 0.060 |
| AAratio:DARPP:zBMI | 0.003 | 1 | 0.955 |
| condition:AAratio:DARPP:zBMI | 8.92 | 3 | 0.030 |

N = 160
Marginal $R^2$ / Conditional $R^2$ = 0.076 / 0.173

**Table S6.** Post hoc effects for the interaction of BMI, condition, and Taq1A

| condition | Taq1A group | estimate | SE | lower 95-CI | higher 95-CL |
|---|---|---|---|---|---|
| ignore | A1- | -0.156 | 0.066 | -0.285 | -0.026 |
| | A1+ | -0.159 | 0.073 | -0.302 | -0.016 |
| **update** | **A1-** | **-0.012** | **0.072** | **-0.153** | **0.129** |
| | A1+ | -0.339 | 0.076 | -0.488 | -0.191 |
| control long | A1- | -0.143 | 0.069 | -0.279 | -0.008 |
| | A1+ | -0.321 | 0.074 | -0.466 | -0.175 |
| control short | A1- | -0.194 | 0.078 | -0.347 | -0.042 |
| | A1+ | -0.260 | 0.081 | -0.421 | -0.099 |

*Note:* the non-significant effect was highlighted in bold.

**Table S7.** Post hoc effects for the interaction of BMI, condition, and DARPP

| condition | DARPP group | estimate | SE | lower 95-CI | higher 95-CL |
|---|---|---|---|---|---|
| ignore | A/A | -0.181 | 0.061 | -0.301 | -0.062 |
| | **G-carrier** | **-0.0625** | **0.076** | **-0.211** | **0.086** |
| update | **A/A** | **-0.044** | **0.066** | **-0.174** | **0.086** |
| | G-carrier | -0.324 | 0.079 | -0.478 | -0.170 |
| control long | A/A | -0.2060 | 0.063 | -0.329 | -0.082 |
| | G-carrier | -0.1689 | 0.078 | -0.322 | -0.016 |
| control short | A/A | -0.194 | 0.069 | -0.331 | -0.057 |
| | G-carrier | -0.213 | 0.086 | -0.380 | -0.045 |

*Note:* non-significant effects are highlighted in bold.

**Table S8.** Full outputs for the models investigating the direct relationship of each SNP and BMI

| | | estimate | SE | t-value | p | adjusted $R^2$ |
|---|---|---|---|---|---|---|
| Taq1A | (intercept) | 26.117 | 0.442 | 59.055 | < .001 | 9.51e-06 |
| | A1+ | 0.742 | 0.741 | 1.002 | 0.317 | |
| COMT | (intercept) | 26.965 | 0.652 | 41.326 | < .001 | -0.001534 |
| | Val/Met | -0.617 | 0.843 | -0.732 | 0.464 | |
| | Met/Met | -1.179 | 0.962 | -1.226 | 0.221 | |
| DARPP-32 | (intercept) | 26.727 | 0.465 | 57.452 | < .001 | 0.0009848 |
| | G-carrier | -0.824 | 0.719 | -1.146 | 0.252 | |
| C957T | (intercept) | 26.307 | 0.714 | 36.844 | < .001 | 0.002643 |
| | A/G | -0.381 | 0.873 | -0.436 | 0.663 | |
| | G/G | 1.080 | 1.007 | 1.073 | 0.284 | |

**Table S9.** Full output for the model investigating the direct relationship of amino acid ratio and BMI

| | estimate | SE | t-value | p |
|---|---|---|---|---|
| (Intercept) | 25.330 | 1.742 | 14.545 | < .001 |
| Amino acid ratio | -7.277 | 7.543 | -0.965 | 0.336 |

N = 160
adjusted $R^2$ = -0.0004356
